# Infection of ratites with clade 2.3.4.4b HPAIV H5N1: Potential implications for zoonotic risk

**DOI:** 10.1101/2025.09.08.674895

**Authors:** Holly A. Coombes, Jacob Terrey, Audra-Lynne Schlachter, Phoebe McCarter, Isabella Regina, Richard Hepple, Natalie McGinn, James Seekings, Jayne Cooper, Benjamin Clifton, Benjamin C. Mollett, Marco Falchieri, Alejandro Nunez, Scott M. Reid, Joe James, Ashley C. Banyard

## Abstract

We detected H5N1 high pathogenicity avian influenza in captive Greater Rhea (*Rhea americana*). Viral genetic analysis revealed the mammalian associated PB2-E627K mutation, indicating selection of mammalian-relevant mutations in ratites. Pathologic investigation of available tissues demonstrated severe multifocal necrotising inflammation, and a strong vasculotropism.

## Background

The global spread of clade 2.3.4.4b high pathogenicity avian influenza viruses (HPAIV) has caused extensive mortality in wild birds and poultry [1]. A defining feature of the H5Nx clade 2.3.4.4b panzootic is its broad host range, characterised by infection across diverse avian species and spillover into multiple mammalian hosts, including humans [2]. An important feature following HPAIV infection of mammals is the adaptation of the viral polymerase proteins, enabling enhanced replication in these hosts. Species-specific adaptations, and their drivers, are rarely defined for these viruses, regardless of the infected host, and a significant number of viral adaptations driven by host-specific factors remain to be elucidated. Here we describe the investigation of viral evolution following infection of rheas and subsequent infection of chickens within a captive environment, where mutational changes in a key viral polymerase protein, PB2, associated with enhanced replication in mammals were detected.

## The Study

An outbreak of H5N1 HPAI was detected in a Greater Rhea (*Rhea americana*) (n = 5) enclosure (Figure 1a) in East Ridings of Yorkshire on 20^th^ December 2024 (Figure 1b). The rheas were housed at a zoological collection of numerous exotic mammals and birds, including chickens (*Gallus gallus domestics*) (Figure 1a). Following onset of clinical disease and subsequent mortality in the rheas, oropharyngeal (OP) and cloacal (C) swabs, along with brain tissue, were collected. Total nucleic acid was extracted from all samples for testing by three AIV real-time reverse transcription polymerase chain reaction (RT-PCR) assays consisting of the Matrix (M)-gene assay for generic influenza A viral RNA (vRNA) detection, an assay for detection of HPAIV H5 vRNA and an N1-specific RT-PCR to detect the neuraminidase (NA) subtype, as previously described [3]. For each assay, samples with a Cq value ≤36.0 were considered positive and sent for whole genome sequencing (WGS).

**Figure 1:**
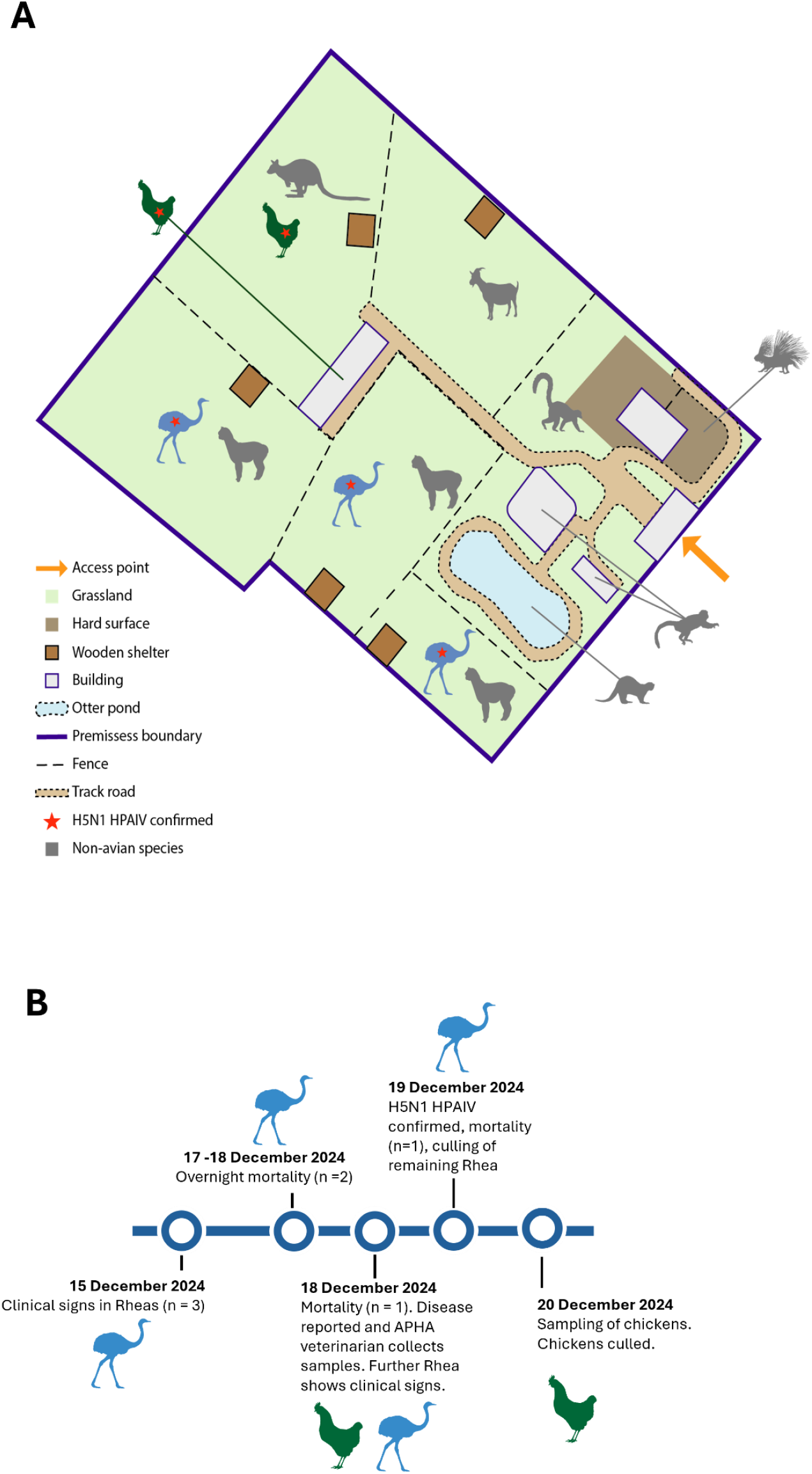
Map of the infected poultry premises with H5N1 HPAIV positive rheas and timeline of key events. (**A**) Map of the infected poultry premises indicating major geographical features and animal location during veterinary visits by APHA. (**B)** A timeline of key events during the infection event.

Following detection of H5N1 HPAIV vRNA in all the sampled rheas (Table 1), OP and C swabs were taken from a group of chickens housed in an enclosure adjacent to the rheas. H5N1 HPAIV vRNA was detected in 54% (n=7/13) of the sampled chickens.

**Table 1.** RT-PCR results and whole genome sequencing on samples collected from Rheas and Chickens housed on the infected premises

| Species | Animal | Sample Type | RT-PCR result |  |  | WGS results |  |  | Strain | Gisaid EPI ID |
| --- | --- | --- | --- | --- | --- | --- | --- | --- | --- | --- |
|  |  |  | (M-gene) | H5HP | N1 | EURL Genotype | PB2-627 | PB2-701 |  |  |
| Rhea | 1 | OP swab | Positive | Positive | Positive | DI.2 | K | D | A/Rhea/England/063177/2024_\|H5N1\|_2024-12-18 | EPI_ISL_20156266 |
|  |  | C swab | Positive | Positive | Positive | DI.2 | K | D | A/Rhea/England/063178/2024_\|H5N1\|_2024-12-18 |  |
|  |  | Brain | Positive | Positive | Positive | DI.2 | K | D | A/Rhea/England/063179/2024_\|H5N1\|_2024-12-18 | EPI_ISL_20156402 |
|  | 2 | OP swab | Positive | Positive | Positive | DI.2 | K | D | A/Rhea/England/063180/2024_\|H5N1\|_2024-12-18 | EPI_ISL_20156403 |
|  |  | C swab | Positive | Positive | Positive | DI.2 | K | D | A/Rhea/England/063181/2024_\|H5N1\|_2024-12-18 | EPI_ISL_20156404 |
|  |  | Brain | Positive | Positive | Positive | DI.2 | K | D | A/Rhea/England/063182/2024_\|H5N1\|_2024-12-18 | EPI_ISL_19659601 |
|  | 3 | OP swab | Positive | Positive | Positive | DI.2 | K | D | A/Rhea/England/063183/2024_\|H5N1\|_2024-12-18 | EPI_ISL_19897983 |
|  |  | C swab | Positive | Positive | Positive | DNS | DNS |  |  |  |
|  |  | Brain | Positive | Positive | Positive | DI.2 | E | D | A/Rhea/England/063185/2024_\|H5N1\|_2024-12-18 | EPI_ISL_20156406 |
|  | 4 | OP swab | Positive | Positive | Positive | DI.2 | K | D | A/Rhea/England/063186/2024_\|H5N1\|_2024-12-18 | EPI_ISL_20156407 |
|  |  | C swab | Positive | Positive | Positive | DI.2 | K | D | A/Rhea/England/063187/2024_\|H5N1\|_2024-12-18 | EPI_ISL_19659622 |
| Chicken | 5 | OP swab | Negative | Negative | Negative |  |  |  |  |  |
|  |  | C swab | Positive | Positive | Positive | DI.2 | K | D | A/Chicken/England/063189/2024_\|H5N1\|_2024-12-18 | EPI_ISL_19897983 |
|  | 6 | OP swab | Negative | Negative | Negative |  |  |  |  |  |
|  |  | C swab | Positive | Positive | Positive | DI.2 | E | D | A/Chicken/England/063191/2024_\|H5N1\|_2024-12-18 | EPI_ISL_19897607 |
|  | 7 | OP swab | Negative | Positive | Positive | DI.2 | K | D | A/Chicken/England/063192/2024_\|H5N1\|_2024-12-18 | EPI_ISL_20156408 |
|  |  | C swab | Negative | Negative | Negative |  |  |  |  |  |
|  | 8 | OP swab | Negative | Negative | Negative |  |  |  |  |  |
|  |  | C swab | Negative | Negative | Negative |  |  |  |  |  |
|  | 9 | OP swab | Negative | Negative | Negative |  |  |  |  |  |
|  |  | C swab | Negative | Negative | Negative |  |  |  |  |  |
|  | 10 | OP swab | Negative | Negative | Negative |  |  |  |  |  |
|  |  | C swab | Negative | Negative | Negative |  |  |  |  |  |
|  | 11 | OP swab | Negative | Negative | Negative |  |  |  |  |  |
|  |  | C swab | Negative | Negative | Negative |  |  |  |  |  |
|  | 12 | OP swab | Negative | Negative | Negative |  |  |  |  |  |
|  |  | C swab | Negative | Negative | Negative |  |  |  |  |  |
|  | 13 | OP swab | Negative | Positive | Negative | DI.2 | K | D | A/Chicken/England/063204/2024_\|H5N1\|_2024-12-18 | EPI_ISL_20156267 |
|  |  | C swab | Negative | Negative | Negative |  |  |  |  |  |
|  | 14 | OP swab | Negative | Negative | Negative |  |  |  |  |  |
|  |  | C swab | Negative | Negative | Negative |  |  |  |  |  |
|  | 15 | OP swab | Negative | **Positive** | Negative | DI.2 | K | D | A/Chicken/England/063208/2024_\|H5N1\|_2024-12-18 | EPI_ISL_20156268 |
|  |  | C swab | Negative | Negative | Negative |  |  |  |  |  |
|  | 16 | OP swab | Negative | Negative | Negative |  |  |  |  |  |
|  |  | C swab | Negative | Negative | Negative |  |  |  |  |  |
|  | 17 | OP swab | **Positive** | **Positive** | **Positive** | DI.2 | K | D | A/Chicken/England/063212/2024_\|H5N1\|_2024-12-18 | EPI_ISL_20156269 |
|  |  | C swab | Negative | Negative | Negative |  |  |  |  |  |
|  | 18 | OP swab | Negative | Negative | Negative |  |  |  |  |  |
|  |  | C swab | Negative | **Positive** | Negative | DI.2 | K | N | A/Chicken/England/063215/2024_\|H5N1\|_2024-12-18 | EPI_ISL_20156409 |

\*OP= oropharyngeal; C=cloacal; \*DNS=did not produce a sequence

WGS were generated from positive samples using Oxford Nanopore Technology (Table 1) as previously described [4]. Assembly, phylogenetic analysis and genotyping of the HPAIV genomes was performed using a custom in-house pipeline publicly available here (<u>APHA-VGBR/WGS_Pipelines/denovoAssembly_ONT_Public.sh</u>). The study-derived sequences were compared against all avian H5 sequences available on GISAID between 1^st^ January 2020 and 28^th^ February 2025. Sequences were genotyped from phylogenetic trees by comparison to known reference sequences for all genotypes currently circulating in the UK. All samples were confirmed as belonging to the EA-2024-DI genotype, falling into the DI.2 sub-genotype, which has been the dominant genotype across Europe during the 2024 – 2025 epizootic year [5].

WGS were assessed for the presence of adaptive mutations that may confer increased replication in mammals. All sequences were aligned on a per segment basis using MAFFT v7.520 [6], manually trimmed to the open reading frame using Aliview version 1.28 [7], translated to amino acids and visually inspected for mutations. All AIV sequences generated in this study are available on GISAID (https://www.gisaid.org, Table 1). Most WGS originating from both rheas and chickens contained 627K in the PB2 gene (PB2-627K) (Table 1). Interestingly, WGS isolated from one of the rheas contained a mixture of PB2-627E, isolated from the brain sample, and PB2-627K, isolated from the OP swab. These sequences represent the first detection of the PB2-627K mutation in the DI genotype in GB.

Three rhea heads were submitted for pathological examination. Grossly, multifocal variably sized raised pale tan plaques were visible on the oropharyngeal and proximal tracheal mucosa in 2/3 birds. Samples of brain, tongue, oropharynx, skin, trachea, eyelid and eye collected for histopathology and histopathologic examination. Immunohistochemistry was performed as previously described [8] using a mouse monoclonal antibody against influenza A viral nucleoprotein (Statens Serum Institute, Denmark). Control tissues and isotypes were assessed alongside to ensure specificity of labelling. Severe multifocal necrotising inflammation was observed in the nasal cavity (3/3), proximal trachea (3/3 birds) (Fig.2), oropharynx (2/3), and eye (3/3), characterised by multifocal to circumferential leukocytoclastic vasculitis, oedema and thrombus formation in the submucosa and necrosis of adjacent structures and in some areas, the overlying epithelium. Abundant co-localised viral antigen was detected in endothelial cells and vascular smooth muscle, and in epithelial cells and macrophages. A mild random multifocal non suppurative encephalitis was identified in 3/3 birds, with viral antigen widely detected in the endothelium of blood vessels within the neuropil and meninges as well as neurons, ependymal cells and the choroid plexus.

**Figure 2:**
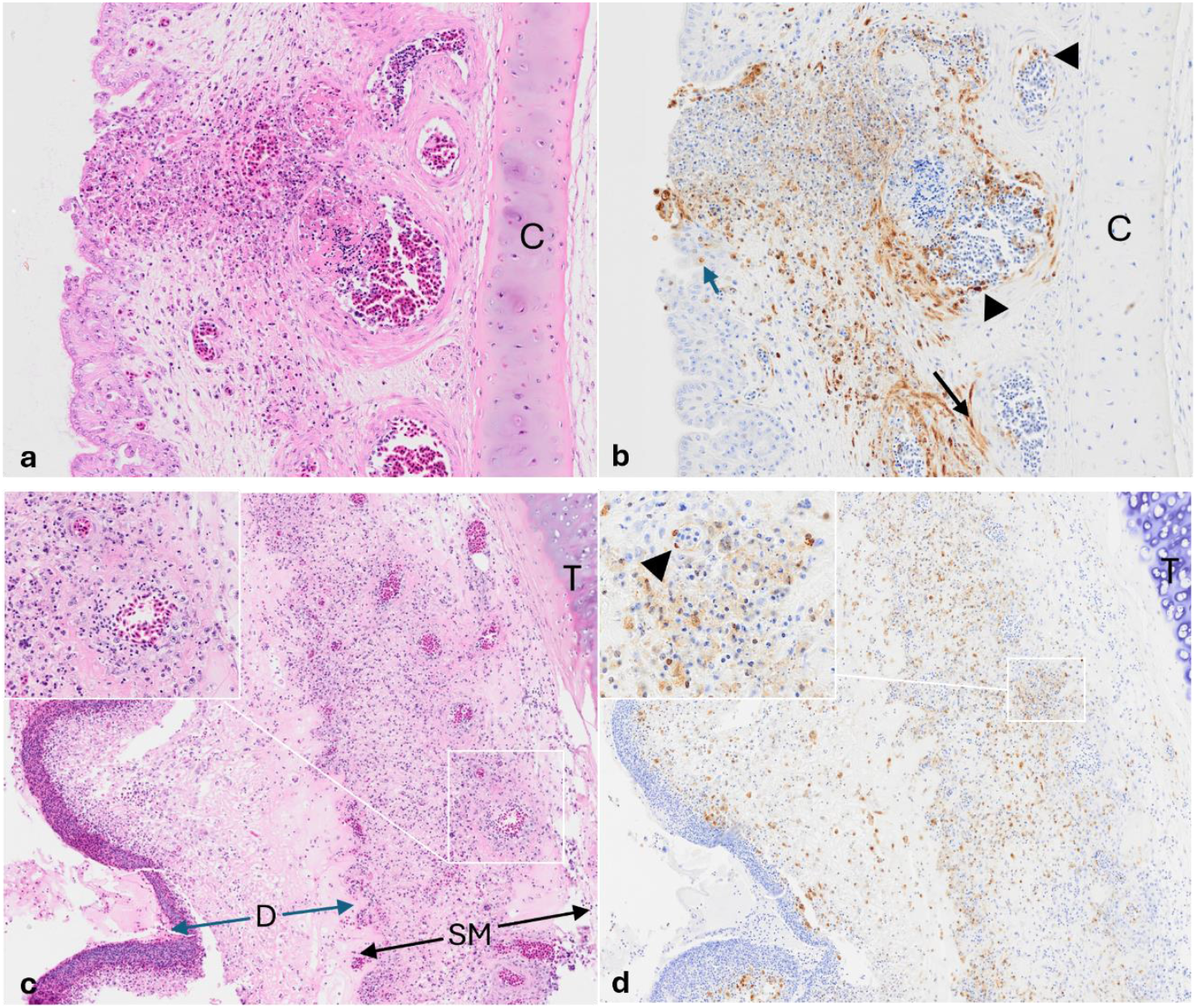
Representative histologic (H&E) and immunohistochemical (IHC) images using an anti influenza A nucleoprotein antibody demonstrating: In the nasal turbinates, a) segmental to circumferential leucocytoclastic vasculitis with thrombus formation in the oedematous submucosa, and necrosis of adjacent structures including overlying epithelium (H&E magnification (mag) x20), with b) co-localised antigen (brown labelling) in the smooth muscle (thin black arrows) and endothelium of capillaries and small vessels (black arrow heads), and in epithelial cells (blue arrow) and macrophages (IHC mag x20). In the trachea, c) similar vascular changes in the submucosa (H&E mag x10, inset mag x40) with necrosis and loss of overlying epithelium, and formation of diphtheritic plaques composed of fibrin, degenerating cells and leucocytes (blue arrows), and d) abundant viral antigen (brown labelling) in endothelial cells (IHC mag x10, inset mag x40, black arrowheads), macrophages and cellular debris. *Abbreviations: C = nasal cartilage, T=trachea cartilage, SM = submucosa, D = diphtheritic membrane*

## Conclusion

To effectively replicate and transmit in mammalian cells, AIVs must overcome multiple host barriers. However, adaptation of AIVs to different avian hosts and variation in host factors is still poorly understood. A key viral adaptation for successful mammalian replication is restored binding of the viral polymerase to host factor acidic nuclear phosphoprotein 32 family member A (ANP32A) [9]. An amino acid change at residue 627 in the PB2 protein, from a glutamate (E) to a lysine (K) is frequently found in mammalian viral sequences [10]. Ratites, along with mammals, lack a 33 amino acid insertion in their ANP32A receptor, typically seen in other avian species, leading to a weaker interaction between the receptor and the viral polymerase [9]. The 627K mutation appears to compensate for this weakened interaction, restoring viral polymerase activity and replication in mammalian cell lines (ref for cell lines) [9]. This may explain why ratites appear to select for 627K mutations, as demonstrated by this study.

Epidemiological and clinical data suggests viral transmission from rheas to co-located chickens, supported by the 627K mutation being present in most of the chicken viral sequences. Although exact transmission chains could not be determined from the sequence data, the persistence of 627K in chickens, despite their avian-like ANP32A, indicates potential maintenance of mammalian adaptive mutations in avian species. Understanding the extent that avian species can maintain mammalian adaptative mutations, is crucial for determine AIV evolution and zoonotic risk. One rhea contained both 627E and 627K viral variants, suggesting within-host viral diversity and viral trophism.

This is the first description of pathologic changes in ratites infected with HPAIV H5N1. Virus induced endothelial damage, vascular inflammation and thrombosis is a known consequence of HPAI H5N1 infection, previously described in cats [11], wild carnivores [12], wild birds [13] and mice [14] in the brain, lungs and eyes. However, the frequent leukocytoclastic inflammation observed in the walls of small to medium vessels, the resultant necrotising inflammation and the abundance of viral antigen detected in most tissues sampled in these birds was unexpected. Viral characteristics (e.g. viral strain, infection route and dose) or host specific factors such as species, age, duration of infection, concomitant disease and immune status likely influenced lesion severity and antigen expression [15]. Despite a limited selection of tissues and small sample size, findings suggest a widespread strong vascular tropism in this species.

Overall, the viral changes detected in the rhea demonstrated early adaptive events following infection of a novel host, including a key viral adaptation that is often associated with increased zoonotic risk. Assessment of viral evolution in microenvironments where unrelated species are co-housed can provide important evidence regarding adaptation to novel hosts.

## Acknowledgements

APHA authors were funded by the Department for Environment, Food and Rural Affairs (Defra, UK) and the Devolved Administrations of Scotland and Wales, through the following programmes: SV3400, SV3032, SE2227 and SE2230. This work was also supported by the Biotechnology and Biological Sciences Research Council (BBSRC) and Department for Environment, Food and Rural Affairs (Defra, UK) research initiative ‘FluTrailMap’ [grant number BB/Y007271/1]. Funded by the European Union under grant agreement (101084171) - (Kappa-Flu). Views and opinions expressed are however those of the author(s) only and do not necessarily reflect those of the European Union or REA. Neither the European Union nor the granting authority can be held responsible for them.

